# Adaptive changes in the fungal cell wall mediate copper homeostasis

**DOI:** 10.1101/2021.12.14.472543

**Authors:** Corinna Probst, Sarela Garcia-Santamarina, Jacob T. Brooks, Inge Van Der Kloet, Dennis J. Thiele, J. Andrew Alspaugh

## Abstract

Copper homeostasis mechanisms are essential for microbial adaption to changing copper levels within the host during infection. In the opportunistic fungal pathogen *Cryptococcus neoformans* (*Cn*), the *Cn* Cbi1/Bim1 protein is a newly identified copper binding and release protein that is highly induced during copper limitation. Recent studies demonstrated that Cbi1 functions in copper uptake through the Ctr1 copper transporter during copper limitation. However, the mechanism of Cbi1 action is unknown. The fungal cell wall is a dynamic structure primarily composed of carbohydrate polymers, such as chitin and chitosan, polymers known to strongly bind copper ions. We demonstrated that Cbi1 depletion affects cell wall integrity and architecture, connecting copper homeostasis with adaptive changes within the fungal cell wall. The *cbi1*Δ mutant strain possesses an aberrant cell wall gene transcriptional signature as well as defects in chitin and chitosan deposition. These changes are reflected in altered macrophage activation and changes in the expression of specific virulence-associated phenotypes. Furthermore, using *Cn* strains defective in chitosan biosynthesis, we demonstrated that cell wall chitosan modulates the ability of the fungal cell to withstand copper stress. In conclusion, our data suggest a dual role for the fungal cell wall, in particular the inner chitin / chitosan layer, in protection against toxic levels of copper and providing a source of metal ion availability during copper starvation. Given the previously described role for Cbi1 in copper uptake, we propose that this copper-binding protein is involved in shuttling copper from the cell wall to the copper transporter Ctr1 for regulated microbial copper uptake.

**Author summary:** Microorganisms must be equipped to readily acquire essential micro-nutrients like copper from nutritionally poor environments while simultaneously shielding themselves from conditions of metal excess. We explored mechanisms of microbial copper homeostasis in the human opportunistic fungal pathogen *Cryptococcus neoformans* (*Cn*) by defining physiological roles of the newly described copper-binding and release protein *Cn* Cbi1/Bim1. Highly induced during copper limitation, Cbi1 has been shown to interact with the high-affinity copper transporter Ctr1. We defined Cbi1-regulated changes in the fungal cell wall, including controlling levels of the structural carbohydrates chitin and chitosan. These polysaccharides are embedded deeply in the cell wall and are known to avidly bind copper. We also defined the host immunological alterations in response to these cell wall changes. Our data suggest a model in which the fungal cell wall, especially the chito-oligomer layer, serves as a copper-binding structure to shield the cell from states of excess copper, while also serving as a copper storage site during conditions of extracellular copper depletion. Given its ability to bind and release copper, the Cbi1 protein likely shuttles copper from the cell wall to copper transporters for regulated copper acquisition.

## Introduction

Metal ions serve important and varied roles in the host-pathogen interaction. Transition metals, such as copper and iron, are essential micronutrients for both the host and pathogen, required as co-factors for cellular respiration and other central cell processes [1, 2]. However, non-bound metal ions can be very cytotoxic. The requirement that microbial cells have ready access to non-toxic levels of transition metals governs a host process called “nutritional immunity”. In this process, the host starves invading microbial pathogens by sequestering essential metals under certain conditions. Conversely, the host may actively bombard the pathogen with toxic levels of metals in other conditions. Nutritional immunity is best studied within microbe-containing macrophages in which Mn, Fe, and Zn are typically restricted by the host, while toxic levels of Cu are actively transported into the phagolysome by the ATP7A copper pump [3].

The genes that control copper homeostasis in fungi are under tight transcriptional control in response to extracellular copper concentrations. In contrast to many other fungi, the human fungal pathogen *Cryptococcus neoformans* (*Cn*) has a single transcription factor, Cuf1, that regulates the transcriptional response to both copper excess and copper starvation [4]. Within the Cuf1 regulon, one of the most highly induced genes during copper starvation is *Cn CBI1/BIM1* (CNAG_02775), encoding a GPI-anchored protein that interacts with the Ctr1 high-affinity Cu^+^ transporter. The *Cn* Cbi1 <u>c</u>opper <u>bi</u>nding and release protein, previously named Bim1, is required for growth in low copper conditions and therefore for effective brain colonization by this neuropathogenic yeast [5]. Cbi1 shares limited homology with lytic polysaccharide monooxygenase (LPMO) proteins that cleave glycosidic bonds within complex carbohydrates such as cellulose, starch, and chitin. Also similar to LPMOs and copper chaperones, *Cn* Cbi1/Bim1 binds copper in a histidine brace region, and it releases copper in reponse to low levels of hydrogen peroxide. However, the purified *Cn* Cbi1/Bim1 protein does not possess the redox activity associated with most sugar-modifying enzymes [6, 7]. It also lacks recognizable polysaccharide binding sites present in LPMOs [6, 7]. Its specific function in Ctr1-mediated uptake of copper is therefore unclear. Based upon previous data, it has been proposed that *Cn* Cbi1/Bim1 acts as an intermediary copper binding protein, delivering copper to Ctr1 for cellular copper acquisition. However, the details of this activity and the source of the copper bound by *Cn* Cbi1/Bim1 are not yet defined [5].

The *Cryptococcus* cell wall is a dynamic structure at the interface between the fungus and its external environment. The basal layer of fungal cell walls is composed primarily of chito-oligomers such as chitin and chitosan, which form a highly cross-linked and rigid structure near the plasma membrane. More superficial layers include other carbohydrates such as α- and β-glucans, as well as mannosylated proteins [8]. During infection and other periods of cell stress, *Cryptococcus* species remodel the cell wall to promote microbial survival under changing environmental conditions. During infection, these adaptive cell wall changes include facilitating the incorporation of the antioxidant pigment melanin, promoting the attachment of an antiphagocytic capsule, and masking immunogenic cell wall epitopes to avoid immune recognition [9-12]. Several host-relevant signals are required for the induction of this type of fungal cell wall remodeling, including host temperature and the relatively alkaline pH encountered during infection [9, 13]. However, it is not well understood how metal stress and nutritional immunity responses affect the fungal cell wall, or if the fungus actively remodels its cell surface in response to those stresses.

Components of fungal cell walls, especially chitin and chitosan, have been previously demonstrated to effectively chelate environmental divalent metal ions such as Cu^2+^ [14, 15] but the physiological importance to the microbe of metal chelation by the fungal cell wall is poorly understood. To further explore the interaction between copper homeostasis and the fungal cell wall, we analyzed the cell wall remodelling response in the *cbi1*Δ mutant strain, which is defective in a cell surface protein involved in copper homeostasis. We first characterized the role of Cbi1 in the transcriptional signatures of *Cn* cell wall-regulating genes. We also defined the physiological effects of mutations in proteins involved in copper homeostasis on the composition and integrity of the cell wall. Since the *Cn* cell wall controls macrophage activation, we determined how copper acquisition and homeostasis affect host innate immune recognition signals as well as the expression of specific microbial phenotypes associated with virulence. These studies suggest that copper availability affects the architecture and integrity of the fungal cell wall; processes likely required for microbial adaptation to host-like nutrional environments. Based upon our data we propose a dual role for the fungal cell wall in protecting *C. neformans* against the presence of toxic levels of copper and providing a source of metal ion availability during copper starvation.

## Results

### The transcription factor Cuf1 regulates cell wall integrity in response to cellular copper levels through the Cbi1-Ctr1 copper uptake complex

A recent study identified the *C. neoformans* Cuf1-dependent copper regulon in response to both copper deficiency and copper excess [4]. During copper deficiency, the most upregulated Cuf1-dependent transcripts represent genes involved in copper uptake, including those encoding for the high-affinity copper transporters Ctr1 and Ctr4 and the newly identified Cbi1 protein [4]. Previously referred to as *Cn* Bim1, this protein has been renamed Cbi1 to reflect its known biochemical activities [6, 7], and to recognize the previously described *Cn* Bim1 microtubule-binding protein involved in filamentous growth [16]. Other *Cn* Cuf1- and copper-regulated genes include many involved in cell wall synthesis and carbohydrate metabolism, potentially connecting copper homeostasis with cell wall remodeling [4, 5].

In pathogens, the fungal cell wall is a dynamic structure required for viability, stress resistance, morphogenesis and virulence. Its composition is actively remodeled in response to various stress signals, and this process is controlled by conserved signaling cascades, including the cell wall integrity (CWI) pathway [17]. The cell wall stress experienced by the copper homeostasis mutants, specifically during copper deprivation, is reflected in transcriptional alterations in the CWI pathway. The *ROM2* gene encodes a guanine nucleotide exchange factor required for CWI pathway activation under conditions of cell stress [18]. Transcriptional induction of *ROM2* is therefore typically observed during conditions of cell wall stress. Comparative transcriptional analysis of the WT and *cuf1*Δ mutant revealed similar *ROM2* transcript levels during copper sufficiency. In contrast, *ROM2* transcript levels were 3-fold higher in the *cuf1*Δ strain compared to wildtype during copper starvation, consistent with an accentuated sensing of cell wall stress in this strain in this condition. Complementation of the *cuf1*Δ mutant with the *CUF1-FLAG* gene (*cuf1*Δ*^C^* strain) restored *ROM2* transcript to wildtype levels (Fig 1B).

**Figure 1:**
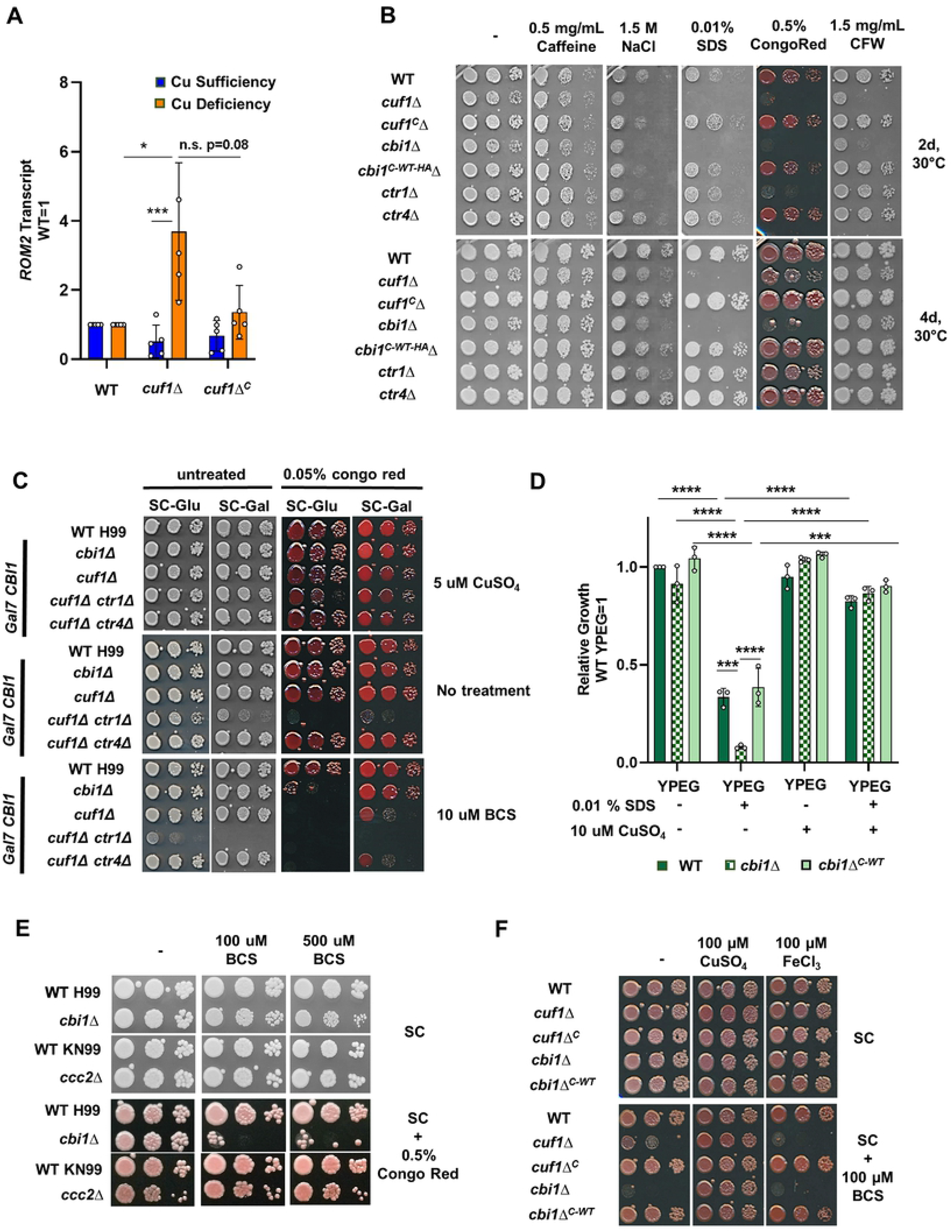
The Cuf1 transcription factor as well as it targets Cbi1 and Ctr1 are involved in maintaining cell wall integrity during copper deficiency. **(A)** qRT-PCR analysis of the *ROM2* transcript level in indicated strains. For the high copper condition, the WT, *cuf1*Δ and Cuf1-Flag complemented *cuf1*Δ*^C^* strains were inoculated to OD_600_ 0.3 in SC supplemented with 1 mM CuSO_4_ and cultivated for 1h at 30°C. To induce low copper conditions, indicated strains were inoculated to OD_600_ 0.3 in SC supplemented with 1 mM BCS and cultivated for 6h at 30°C. For comparison the WT *ROM2* transcript levels at each condition were set to 1. Presented is the mean +/-SEM of the relative transcript levels of 4 biological replicates. A 2-way ANOVA was performed using GraphPad Prism from log transformed data. **(B)** Growth analysis in presence of cell wall/ surface stressors. The spotting assay was performed on SC supplemented with 100 µM BCS which was co-supplemented with indicated amounts of cell wall and cell surface stressors. Indicated strains were grown overnight in SC at 30°C. Cells were diluted to OD_600_ 0.25 and a serial 1:10 dilution was spotted onto media plates. Plates were incubated at 30°C for 2-4d. This figure shows a representative image from 3 independent spotting experiments. **(C)** *GAL7* promoter-driven expression of Cbi1 in different *cuf1*Δ *ctr*Δ strains during cell wall stress. The spotting assay was performed on SC with either glucose (SC-Glu) or galactose (SC-Gal) as carbohydrate source, supplemented with indicated amounts of CuSO_4_, BCS and congo red. Indicated strains were grown overnight in SC at 30°C. Cells were diluted in PBS to OD_600_ 0.25, and a serial 1:10 dilution was spotted on to media plates. Plates were incubated at 30°C for 3d. This figure shows a representative image from 3 independent spotting experiments. (D) Growth of WT, *cbi1*Δ and Cbi1 WT complemented *cbi1*Δ (*cbi1*Δ*^C-WT^*) in presence of 0.01% SDS in YPEG w/o 5 µM CuSO_4_. Growth was measured via OD_600_ after 72h at 30°C and was normalized to WT growth in YPEG media. Presented is the mean +/-SEM of the relative growth rates of 3 biological replicates. 1-way ANOVA was performed using GraphPad Prism from log transformed data. **(E)** Impact of the *ccc2*Δ mutation on growth in presence of the cell wall stressor Congo red. The spotting assay was performed on SC supplemented with 100 µM or 500 μM BCS which was co-supplemented with 0.5% Congo red. Indicated strains were grown overnight in SC at 30°C. Cells were diluted to OD_600_ of 0.25 and a serial 1:10 dilution was spotted on to media plates. Plates were incubated at 30°C for 2-4d. (D) Copper restoration of cell wall stressor sensitivity of *cbi1*Δ cells. The spotting assay was performed on SC+0.5 % congo red supplemented with indicated amounts of BCS, CuSO_4_ and/or FeCl_3_. Indicated strains were grown overnight in SC at 30°C. Cells were diluted to OD_600_ of 2.5 and a serial 1:10 dilution was spotted on to media plates. Plates were incubated at 30°C for 3d. This figure shows a representative image from 3 independent spotting experiments.

To further analyze the impact of *CUF1* and other copper homeostasis genes for maintaining cell wall integrity during copper deficiency, we assessed the sensitivity of the *cuf1*Δ, *cbi1*Δ, *ctr1*Δ and *ctr4*Δ strains to cell wall stressors in copper-replete and copper-deficient conditions. These stresses included calcofluor white (CFW, blocks chitin assembly), Congo red (impairs assembly of cell wall polymers, mainly chitin), caffeine (impacts PKA-mediated signal transduction), SDS (cell surface/ membrane stressor) and NaCl (osmotic stressor) [19-21]. We did not observe any growth phenotypes for the mutant strains in the presence of these cell wall stressors under copper-replete conditions (Supp Fig 1A). In contrast, during copper starvation induced by the extracellular copper (I) chelator bathocuproinedisulfonic acid (BCS) (100μM), the *cuf1*Δ *and cbi1*Δ strains exhibited strong growth inhibition compared to WT on NaCl, SDS, Congo red and CFW-containing media (Fig 1B). Even at more modest levels of copper chelation (10μM BCS), the *cuf1*Δ *and cbi1*Δ strains exhibited strong growth inhibition on Congo red-containing media (Fig. 1C). Additionally, the Ctr1 copper transporter was similarly required for survival during copper deprivation in the presence of NaCl and CFW, but not in the presence of SDS. No cell wall stress-associated growth defect was noted for the *ctr4*Δ copper transporter mutant strain (Fig 1B), supporting prior observations that the Ctr1 and Ctr4 high affinity copper transporters serve overlapping but non-redundant functions in *Cn* copper homeostasis [22]. Growth inhibition of the *cuf1*Δ and *cbi1*Δ strains was complemented by expressing epitope-tagged versions of these proteins in the respective mutant strains [Cuf1-Flag in *cuf1*Δ (cuf1Δ^C^) or Cbi1-HA in *cbi1*Δ (*cbi1*Δ^C-WT-HA^)]. Together, these results suggest that copper homestasis is required for *Cn* cell wall stress resistance.

To further explore the relationships among the *Cn* copper homeostasis proteins in cell wall integrity, especially the less well-characterized Cbi1 protein, we conditionally overexpressed the *CBI1* gene in a series of individual and double mutant strains (Fig 1C). Galactose-mediated overexpression of the p*GAL7-CBI1* allele fully complemented the copper-dependent cell wall growth defect of the *cbi1*Δ mutant, and partially that of the *cuf1*Δ strain. In contrast, overexpression of *CBI1* was unable to restore cell wall integrity to strains with a *ctr1*Δ mutation. These results are consistent with prior studies suggesting that Cbi1 and Ctr1 are independent components of a copper transporter complex [5]. Additionally, these findings indicate that defective Cbi1 function is likely responsible for much of the loss of cell wall integrity in the *cuf1*Δ mutant.

We also assessed *Cn* copper-dependent cell wall integrity phenotypes using an alternative method of copper limitation to extracellular copper chelation by BCS. We incubated the *Cn* strains in media containing ethanol and glycerol as sole carbon sources. Growth on these non-fermentable carbohydrates is only supported by cellular respiration, effectively shunting intracellular copper into the mitochondria to the copper-dependent enzymes required for oxidative phosphorylation. Incubation of the WT and *cbi1*Δ*^C-WT^* strains in YPEG + 0.01% SDS caused a ∼40% growth reduction, and an even more severe reduction of growth (∼90%) in the *cbi1*Δ strain (Fig 1D). This growth impairment was complemented in all strains by supplementation with CuSO_4_, suggesting that shuttling of copper from other pathways towards respiration influences the ability of *Cn* to withstand cell surface stress. Depletion of Cbi1 further decreased cell fitness under these conditions.

We also tested the cell wall integrity of the *Cn ccc2*Δ mutant, a strain defective in copper transport withinin the secretory pathway and subsequent altered metalation of secreted proteins [2, 23]. A modest growth defect on Congo red was observed for the *ccc2*Δ strain, and the defect developed independent of copper availability (Fig. 1E). These results suggest that defective copper loading of enzymes in the secretory compartment is likely not the cause of the *Cn* cell wall phenotypes observed during copper deficiency.

The Cfo1 ferroxidase is a copper-dependent enzyme involved in high-affinity iron acquisition [24]. Therefore, loss-of-function mutations in Cuf1 or other components of the *Cn* copper uptake machinery would be predicted to affect intracellular iron concentrations as well as copper levels. Additionally, previous studies demonstrated that iron homeostasis is important for proper fungal cell wall and membrane architecture [25, 26]. We therefore analyzed the effects of exogenous copper or iron on the BCS-induced cell wall phenotypes of the *cbi1*Δ and *cuf1*Δ mutants. Individual supplementation of the growth medium with copper,but not iron, restored growth to the *cbi1Δ* and *cuf1Δ* strains in the presence of cell wall stress and copper depletion (Fig 1F).

### Changes in *Cn* cell wall composition in response to defective copper homeostasis

To further characterize the specific role of Cbi1 in cell wall homeostasis during copper stress, we used transmission electron microscopy (TEM) to characterize the cell wall architecture of the wildtype (WT) and *cbi1*Δ strains incubated in both copper-sufficient and copper-deficient growth conditions (Fig 2). In copper-sufficient conditions the *Cn* WT cell wall consists of two layers characterized by differing electron density [27-29]. Extracellular copper sequestration by the highly copper-specific chelator BCS resulted in decreased electron density in the innermost cell wall layer composed primarily of chitin and chitosan (Fig 2 A, C). These chito-oligomers efficiently bind bivalent metals (including copper ions), consistent with the higher electron density of this cell wall layer during copper sufficiency [14, 15]. The cell wall of the *cbi1*Δ mutant strain was similar to WT during copper sufficiency, displaying distinct layers based on electron density. However, the copper-starved *cbi1*Δ strain demonstrated a reduction in total cell wall thickness compared to both the WT strain and the copper-sufficient *cbi1*Δ cells (Fig 2B). Also in contrast to WT, there was no reduction in the inner cell wall electron density in the *cbi1*Δ mutant strains during copper deficiency (Fig 2A, C). These results suggest a model in which Cbi1 is required for the release of cell wall-bound metals during copper starvation.

**Figure 2:**
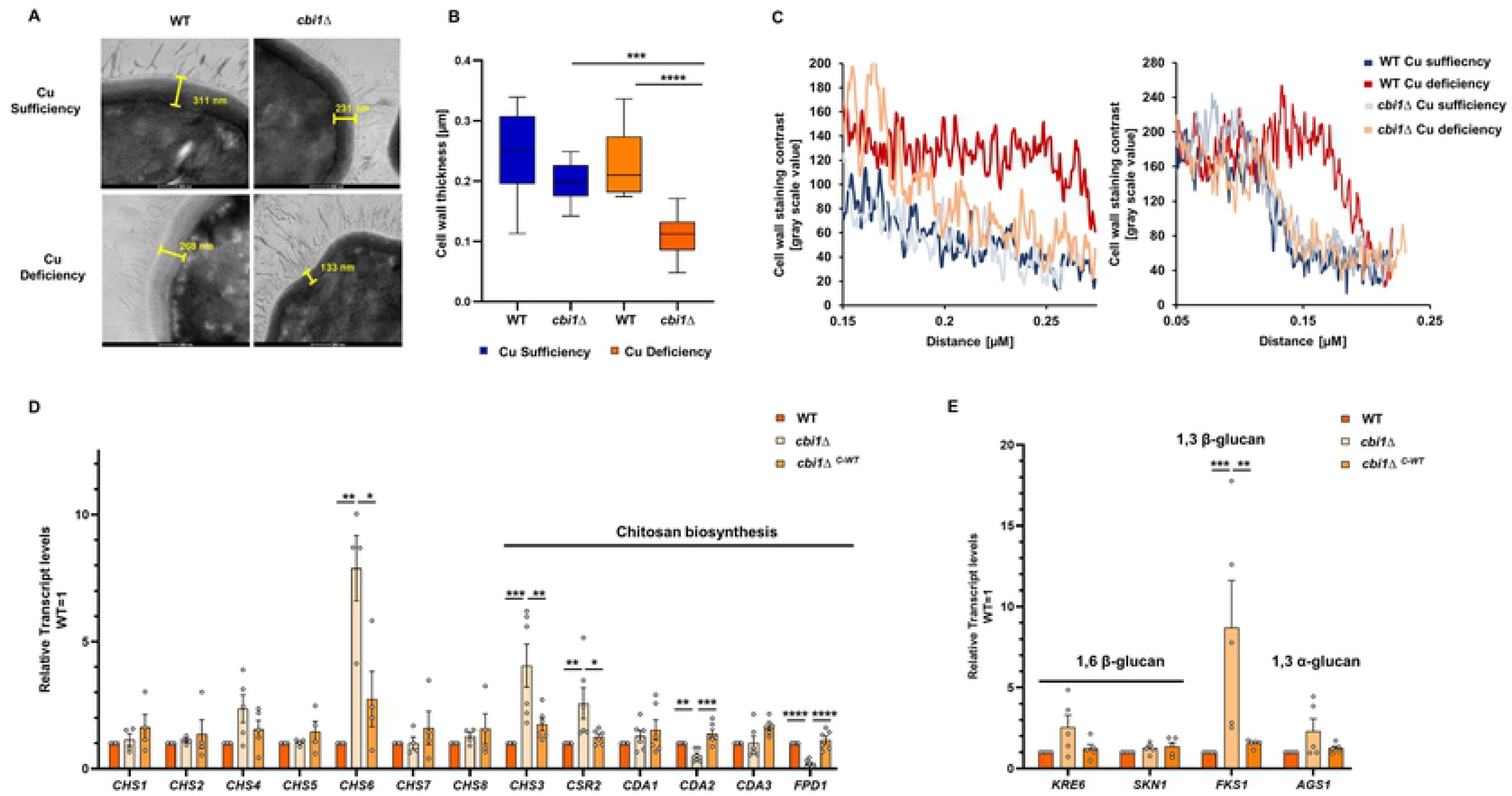
Copper deficient *cbi1Δ* cells show a significant loss of cell wall carbohydrates, reflected by transcriptional mis-regulation in cell wall-associated genes. TEM images of copper sufficient or deficient WT and *cbi1Δ* cells. Cells were incubated in YPD supplemented with 10 µM CuSO_4_ (Cu sufficiency) or with 250 µM BCS (Cu deficiency) for 24h at 30°C. **(A)** Representative images of the TEM analysis (in 29000 x magnification). **(B)** Quantification of cell wall thickness. Measurements were performed using the ImageJ/Fiji measurement tool: Cu sufficiency-WT 10 cells, Cu sufficiency-*cbi1*Δ 8 cells, Cu deficiency-WT 11 cells and Cu deficiency-*cbi1*Δ 14 cells. A 1-way ANOVA was performed using GraphPad Prism from log transformed data. **(C)** Quantification of the cell wall staining intensity. Presented is the analysis of two sets of TEM images. The gray value was measured with ImageJ/Fiji and plotted against the distance along the cell wall. **(D)** qRT-PCR analysis Transcripts involved in chitin and chitosan biosynthesis in copper-deficient WT, *cbi1*Δ, and complemented *cbi1*Δ (*cbi1*Δ*^C-WT^*) cells. Cells were inoculated to OD_600_ 0.05 in YPD supplemented with 250µM BCS and cultivated for 24h at 30°C. For comparison, the WT transcript levels were set to 1. Presented is the mean +/-SEM of the relative transcript levels of minimum 4 biological replicates. A 1-way ANOVA was performed using GraphPad Prism from log transformed data. **(E)** qRT-PCR analysis Transcripts involved in glucan biosynthesis in copper deficiency. Cells were treated as described in (A). Presented is the mean +/-SEM of the relative transcript levels of minimum 4 biological replicates. A 1-way ANOVA was performed using GraphPad Prism from log transformed data.

The ultrastructural changes observed in the *cbi1*Δ strain cell wall during copper starvation were not due to altered cell viability. Although the *cbi1*Δ strain demonstrated a defect in proliferation during copper deficiency, we observed no decrease in *cbi1*Δ cell viability during the first 24 hours, as assessed by quantitative cultures of colony-forming units (CFUs) (Supp Fig 1 B-C. We also assessed the effective “copper state” of the WT and *cbi1*Δ strains in the conditions chosen for copper sufficiency (10μM CuSO_4_) and deficiency (250μM BCS) by quantifying transcript levels of the *CMT1* metallothionein gene (induced during high copper states) and the *CTR4* copper transporter gene (induced during copper starvation) (Supp Fig 1D). Both the WT and *cbi1Δ* strains demonstrated a strong induction of *CTR4* expression, but not of the *CMT1* transcript, indicating that copper deficiency was induced by BCS treatment compared to the copper sufficient growth conditions. We also performed ICP-MS-based metal quantification of cell-associated copper and iron in each growth condition (Supp Fig 1 E-F). A consistent and similar pattern of reduced cell-associated Cu was measured in the WT and *cbi1*Δ cells cells incubated in BCS. Additionally, a decrease in iron levels was observed in copper-deficient cells, consistent with known copper-dependent iron uptake mechanisms in *Cn*.

### Transcriptional changes in cell wall genes in response to copper status

Prior investigations have established a set of *C. neoformans* genes encoding enzymes involved in the synthesis and chemical modification of the major cell wall structural carbohydrates. These include 8 chitin synthetase genes (*CHS1-8*), 4 chitin deacetylase genes (chitin-to-chitosan conversion) (*CDA1-3*, *FPD1*), and the chitin synthase regulator-2 (*CSR2*). Genes involved in glucan synthesis include *KRE6* and *SKN1* (β-1,6-glucan), *FKS1* (β-1,3-glucan) and *AGS1* (α-1,3-glucan) [8]. To investigate how copper availability and Cbi1 might affect cell wall carbohydrate homeostasis, we measured transcript levels of these major cell wall synthesis genes in copper deficient wildtype (WT), *cbi1*Δ mutant, and *cbi1Δ^C-WT^* complemented strain (Fig 2 D-E). Among the genes involved in chitin and chitosan synthesis, we observed significant increases in transcript abundance in the *cbi1*Δ mutant for *CHS6* (∼7 fold), *CHS3* (∼4 fold) and *CSR2* (∼3 fold), while the chitin/chitosan deacetylase genes *CDA2* and *FPD1* were downregulated (Fig 2D). In addition to genes involved in chitin/chitosan biosynthesis, the transcript level of the β-1,3-glucan synthase *FKS1* gene was induced 8-fold. Changes in the expression of *KRE6* (β-1,6-glucan synthesis) and *AGS1* (α-1,3-glucan synthesis) were not statistically significantly altered (Fig 2E). Taken together, these findings indicate that the copper-deficient *cbi1*Δ strain displays transcriptional changes in several cell wall polysaccharide synthesis genes, especially those associated with chitosan and β-1,3-glucan synthesis.

To explore the functional relevance of changes in *Cn* cell wall gene transcript abundance as a function of copper availability, we quantified β-glucan and chitin/chitosan levels in the WT, *cbi1Δ* mutant, and *cbi1Δ^C-WT^* complemented strains after incubation for 24h in copper-sufficient (YPD + 10 µM CuSO_4_) and copper-deficient (YPD + 250 µM BCS) conditions. No significant changes were detected in total cell wall β-glucan between the WT and *cbi1*Δ cells in either growth condition (Supp Fig 2A). However, the *cbi1*Δ mutant exposed to copper-deficiency displayed a greater than 50% reduction in total cell wall chitin and in chitosan compared to WT and complemented strains (Fig 3 A-B). These results are consistent with reduced transcript levels for the *CDA2* and *FPD1* chitin deacetylase genes in the copper-starved *cbi1*Δ strain. Therefore, the reduction of cell wall thickness and altered cell wall integrity in the *cbi1*Δ strain during copper starvation is, in part, likely due to a reduction in the inner cell wall chito-oligomer layer.

**Figure 3:**
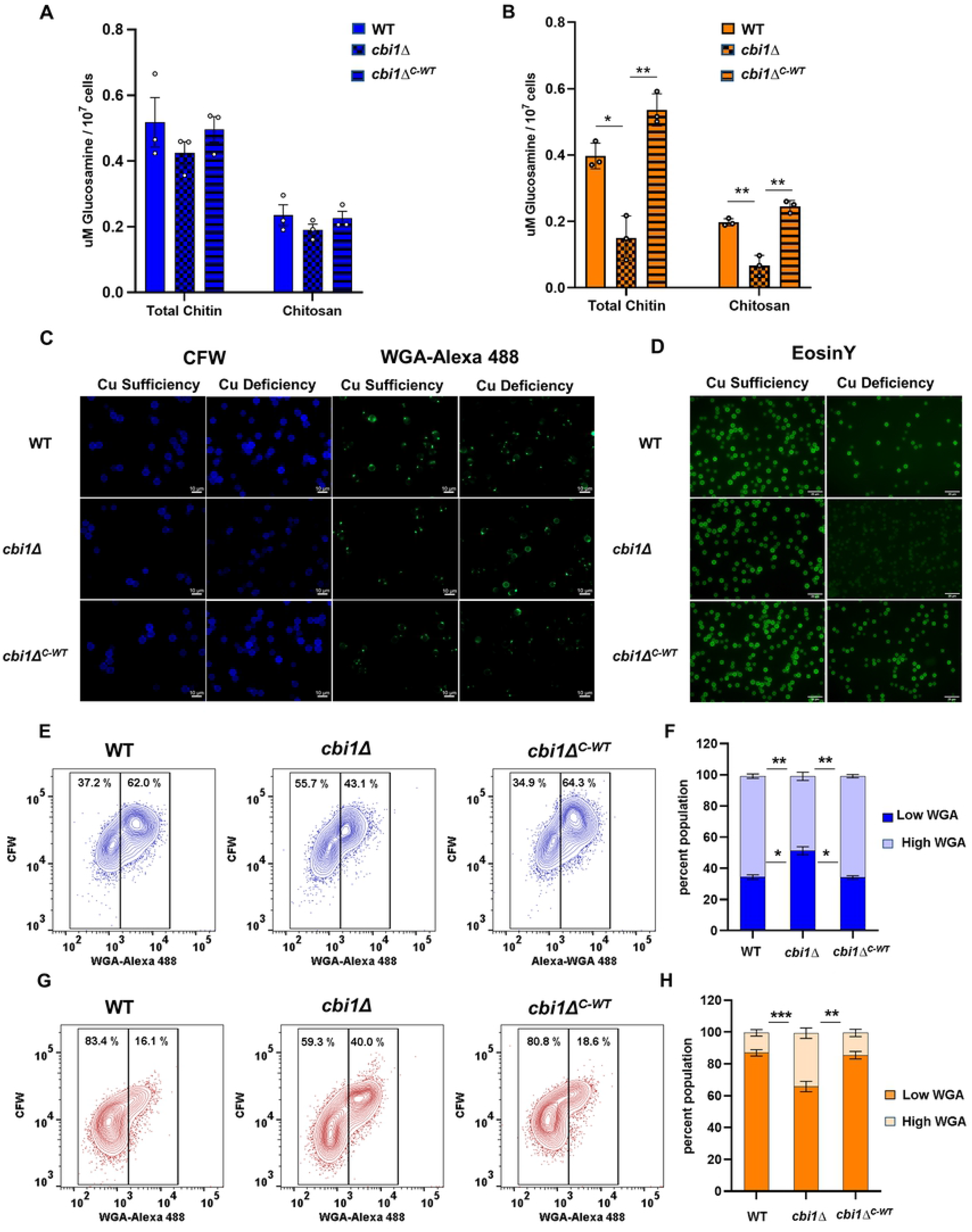
Cbi1 affects cell wall chitin and chitosan deposition and architecture during copper deficiency. **(A)-(B)** MBTH based chitin/chitosan quantification from purified cell wall material of copper sufficient **(A)** or deficient **(B)** WT, *cbi1*Δ and Cbi1 WT complemented *cbi1Δ* (*cbi1*Δ*^C-^*^WT^) cells. Strains were incubated for 24h in YPD+ 10 uM CuSO4 (copper sufficiency) or YPD +250 uM BCS (copper deficiency). Values are shown in uM Glucosamin/10^7^ cells. Presented is the mean +/-SEM of 3 biological replicates. A 1-way ANOVA was performed using GraphPad Prism from log transformed data. **(C)** Calcofluor white (CFW) and wheat germ agglutin (WGA)-Alexa 488 staining for chitin of copper sufficient or deficient WT, *cbi1*Δ and and Cbi1 WT complemented *cbi1*Δ (*cbi1*Δ*^C-^*^WT^) cells. Strains were as described in (A) and double stained with CFW and WGA-Alexa 488. Shown are representative images. Three independent treatments and stainings were performed. **(D)** EosinY staining for chitosan of copper sufficient and deficient WT, *cbi1*Δ and Cbi1 WT complemented *cbi1*Δ (*cbi1*Δ*^C-^*^WT^) cells. Strains were cultivated as described in (A), followed by EosinY staining. Shown are representative images. Five independent treatments and staining were performed. **(E-H)** FACS analysis of CFW and WGA-Alexa 488 stained cells. WT, *cbi1*Δ and *cbi1*Δ*^C-WT^* complemented cells were incubated for 24h in YPD+ 10 uM CuSO4 (E, F; copper sufficiency) or YPD +250 µM BCS (G,H; copper deficiency), harvest and double stained with CFW and WGA-Alexa 488. Stained cells were analyzed using a FACS Canto A Analyzer and data were analyzed using Flow Jo. **(E)** and **(G)** show representative FACS profiles of indicated copper sufficient or deficient strains, **(F)** and **(H)** show the quantification of the population pattern from 3 independent experiments. A 2-way ANOVA was performed using GraphPad Prism from log transformed data.

To examine detailed changes in patterns of chitin and chitosan deposition, we performed microscopy using chitin- and chitosan-specific fluorescent stains. We double-stained Cu-sufficient and Cu- deficient WT, *cbi1*Δ and *cbi1*Δ*^C-WT^* cells with Calcofluor white (CFW), a small molecule globally staining chitin, and AlexaFluor488-conjugated wheat germ agglutin (WGA-Alexa 488), a lectin that binds exposed chito-oligomers. To a similar extent as in the biochemical chitin assays, we observed a reduction in CFW staining intensity of Cu-deficient *cbi1Δ* cells (Fig 3 C, Supp Fig 2B). WGA staining of WT and *cbi1*Δ*^C-WT^* complemented cells only revealed exposed chito-oligomers, primarily at regions of cell separation, budding sites, and bud scars (Fig 3 C). In contrast, the copper-deficient *cbi1Δ* cells demonstrated an enrichment of WGA-Alexa 488 staining globally around the cell surface. We additionally stained cells with EosinY, a small molecule that binds to chitosan polymers. Consistent with the biochemical data indicating reduced chitosan levels, we observed decreased EosinY staining intensity in copper-starved *cbi1*Δ cells (Fig 3D, Supp Fig 2 C-D).

We also performed flow cytometry on CFW and WGA-Alexa 488 stained cells to more precisely quantify the altered chito-oligomer staining pattern in these strains (Fig 3 E-H, Supp Fig 2 E-F). We directly compared WGA-Alexa 488 staining intensity to CFW staining intensity to assess chito-oligomer exposure relative to total cell wall chito-oligomer content. In copper-sufficient growth conditions, there was a small but statistically significant decrease in WGA-Alexa 488 staining intensity among the *cbi1*Δ cells compared to WT and complemented strains. However, we observed a notable increase in relative WGA staining in a sizable subpopulation of *cbi1*Δ cells (“High WGA”, Fig 3 G-H, Supp Fig 2E) during copper-starvation, consistent with the fluorescent microscopy results. Together, these findings suggest that the cell wall chito-oligomers of copper-deficient *cbi1Δ* cells are not only decreased in total amount, but they are deposited within the cell wall in an aberrant manner, leading to a higher degree of exposure.

To explore the role of chitin and chitosan for modulating the resistance to copper stress, we assessed the growth effects during low and high copper stress for strains with mutations in either copper homeostasis or chitosan synthesis (Fig 4). As previously described, we observed poor growth of the *cbi1*Δ and *ctr1*Δ strains during copper deficiency (Fig 4B). In line with previous findings, no sensitivity to copper starvation was observed for the *ctr4*Δ copper transporter mutant strain [30]. These results indicate that the Cbi1/Ctr1 complex can likely compensate for the loss of Ctr4, but that the Ctr4 transporter is not sufficient to maintain *Cn* growth during copper starvation in the absence of Ctr1. A slight reduction in growth was observed for the *chs3*Δ strain during copper starvation. Since Chs3 is responsible for the synthesis of most of the chitin destined to be converted to chitosan, this result suggests that reductions in chitosan may affect the ability of *Cn to* withstand low copper stress. No growth phenotype was observed for the *cda2*Δ chitin deacetylase gene, indicating that the observed dysregulation of *CDA2* in the *cbi1*Δ background does not explain the *cbi1*Δ growth defect during copper deficiency.

**Figure 4:**
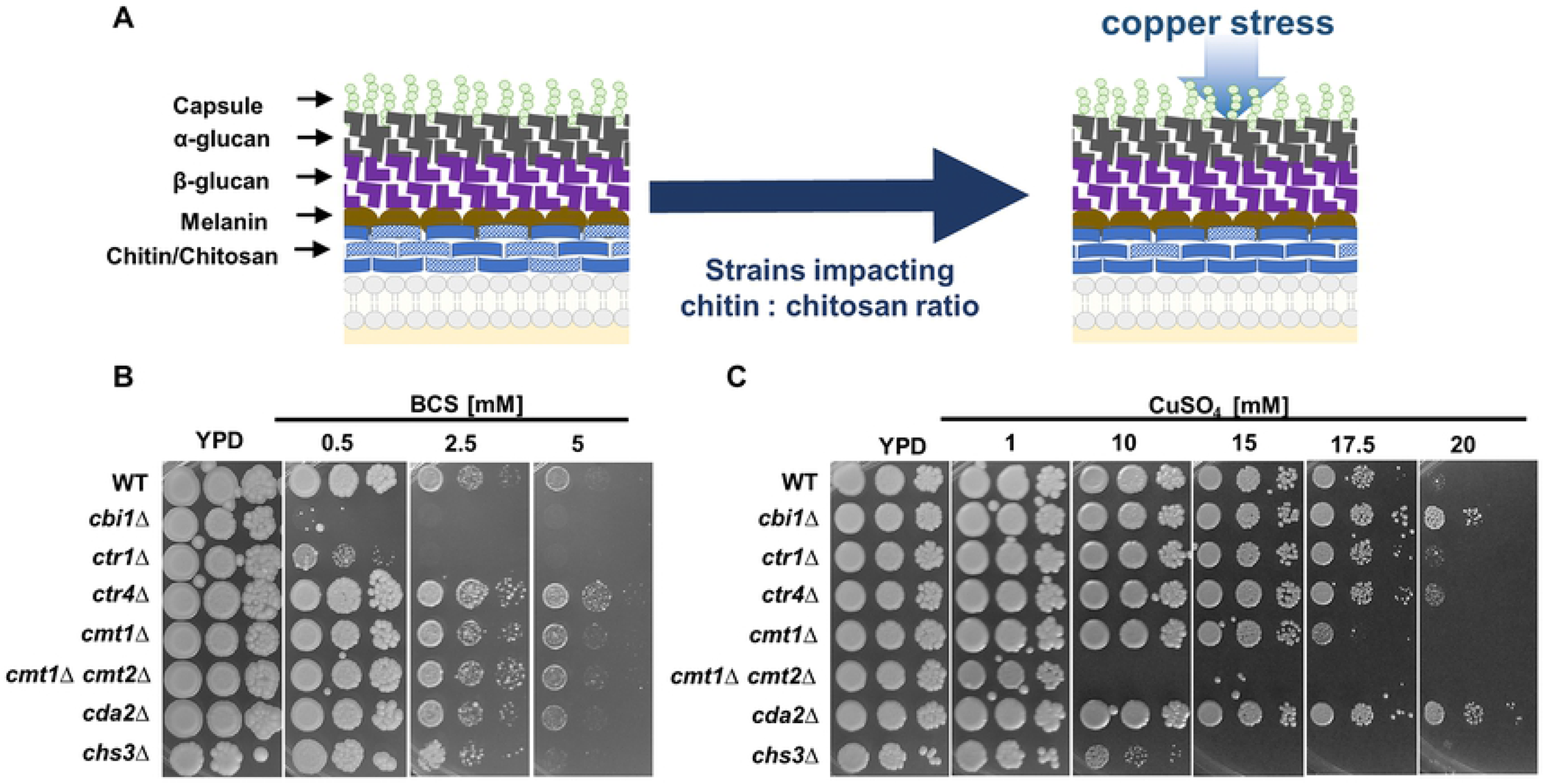
Chitosan deposition in the cell wall influences *C. neoformans* growth during copper stress. **(A)** Proposed model for the role of chitosan deposition to withstand copper stress. The cryptococcal cell wall is a dynamic multi-layered compartment build up by the sugar polymers chitin, α- and β-glucans. Chitin (solid blue) and its deacetylated form chitosan (dotted blue) build the inner cell wall layer, which is covered by an upper layer of β-glucans (purple) and α-glucans (black). The pigment melanin (brown circles) is incorporated and attached to cell wall chitosan, and the cryptococcal capsule (green dots) is attached to α-glucans. **(B-C)** Growth analysis in the presence of low (B) and high (C) copper stress. The spotting assay was performed on YPD supplemented with indicated amounts of BCS or CuSO_4_. Indicated strains were grown overnight in YPD at 30°C. Cells were diluted to OD_600_ of 0.2 and a serial 1:10 dilution was spotted on to media plates. Plates were incubated at 30°C for 2-6d.

To assess the role of the chitin and chitosan inner layer on resistance to copper toxicity, we tested the *chs3*Δ and *cda2*Δ mutant strains for growth phenotypes in the presence of increasing copper concentations. The chitosan-deficient *chs3Δ* strain was more sensitive to high copper stress than the wild-type, and similar in its copper sensivity to strains with mutations in the *Cn* metallothionein genes *CMT1* and *CMT2* that mediate scavenging of excess copper (Fig 4C). In contrast to *chs3*Δ, the *cda2*Δ strain, with a mutation in a single chitin deacetylase gene but relatively preserved cell wall chitosan levels, demonstrated resistance to toxic copper levels compared to WT. The *cbi1*Δ mutant displayed a similar copper resistance profile as the *cda2*Δ strain. This increased copper resistance was not shared with the *ctr1*Δ or *ctr4*Δ copper transporter mutants, suggesting that altered copper transport was not responsible for this phenotype. This finding suggests an unexpected new role for Cbi1 in modulating growth during high copper stress, potentially by its known role in modulating *CDA2* function.

### Cell wall changes in response to low copper stress lead to increased macrophage activation and altered caspofungin tolerance

We tested the physiological relevance of altered copper homestasis to other infection-related processes involving the cell wall. *C. neoformans* strains with enhanced exposure of cell wall chito-oligomers often display increased activation of host innate immune cell activity [31, 32]. To investigate the physiological consequences of the aberrant *cbi1*Δ cell wall structure in host cell interactions, we co-incubated *C. neoformans* strains with murine bone marrow-derived macrophages (BMM), assessing TNF-α production as a marker of host immune cell activation. Macrophages exposed to copper-deficient *cbi1*Δ cells showed a statistically significant increase in TNF-α secretion compared to macrophages co-incubated with similarly treated WT or complemented strains (Fig 5A). No differences in TNF-α production were noted for macrophages incubated with any of these strains grown in the presence of copper.

**Figure 5:**
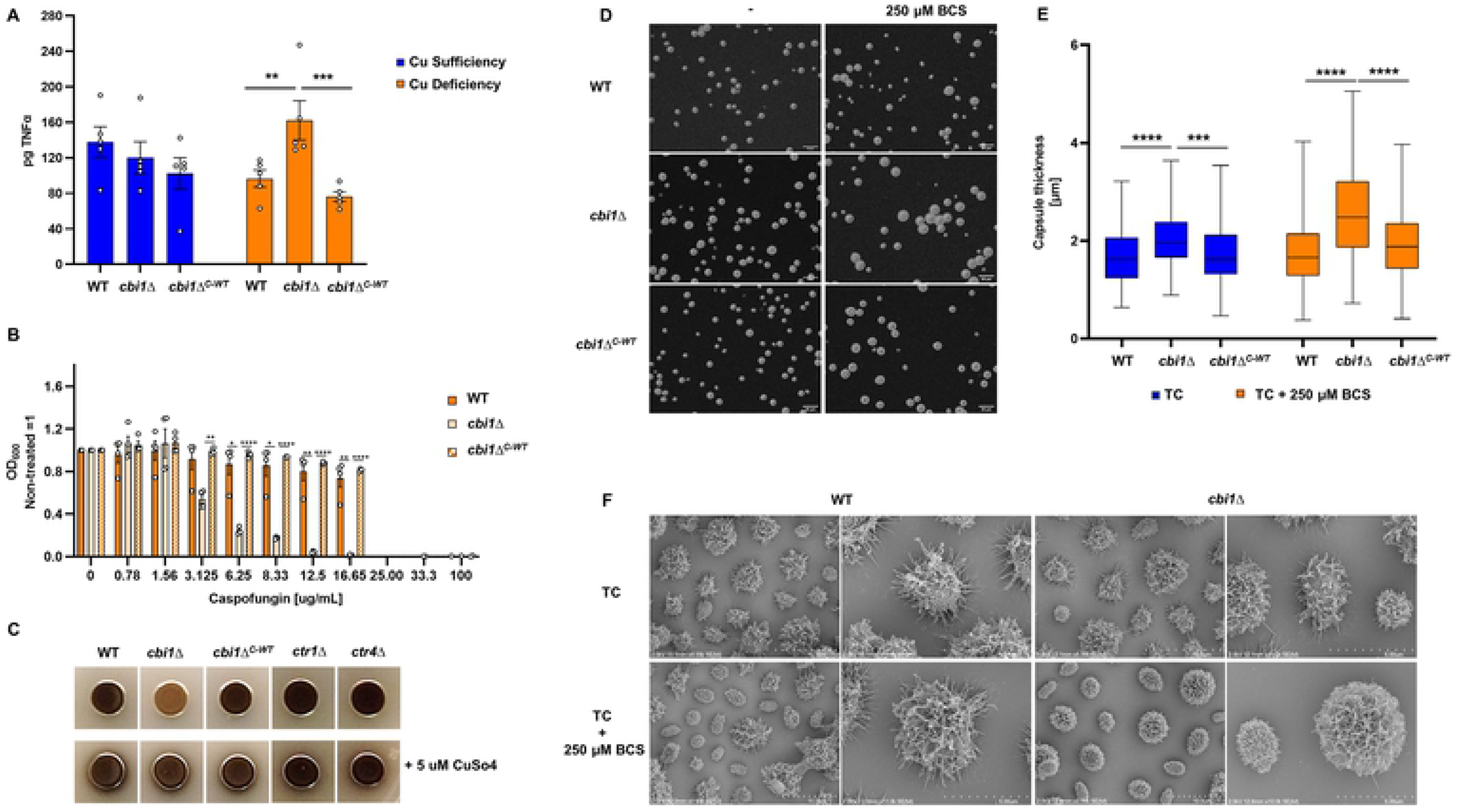
Cbi1 depletion and copper deficiency impacts several cell wall associated virulence phenotypes. **(A)** Macrophage activation assay upon infection with copper sufficient or deficient WT, *cbi1*Δ and *cbi1*Δ*^C-WT^* complemented cells. BMMs were harvested from A/J mice and co-incubated with *Cn* cells at an MOI of 10:1 (Cn:BMMs), followed by an ELISA-based quantification of TNF-α (pg) in the supernatant. Presented is the mean +/-SEM of 5 independent experiments. A 2-way ANOVA was performed using GraphPad Prism from log transformed data. **(B)** Minimal inhibitory concentration (MIC) analysis of caspofungin during copper deficiency. WT, *cbi1*Δ and *cbi1*Δ*^C-WT^* were grown in 96-well liquid cultures in SC media supplemented with 100 μM BCS and were co-treated with 0 to 100 ug/mL caspofungin. OD_600_ was measured after 24h of growth at 30°C and the OD_600_ of the non-treated condition was set to 1. Presented is the mean +/-SEM of 4 independent experiments. A 2-way ANOVA was performed using GraphPad Prism from log transformed data. **(C)** Melanization of WT, *cbi1*Δ, *ctr1*Δ and *ctr4*Δ cells in the absence and presence of 5 µM CuSO_4_. Overnight cultures were harvested, washed 1x with PBS, diluted to OD_600_ 2.5 and spotted on to L-DOPA plates. Shown are representative images from 3 independent experiments. **(D)** Analysis of capsule formation using India ink contrast staining of WT, *cbi1*Δ and *cbi1*Δ*^C-WT^* complemented cells. Indicated strains were grown for 3d in capsule inducing conditions in presence and absence of 250 µM BCS. Cell were harvest, resuspended in PBS, stained with India ink (1:1) and analyzed using the DIC channel. Shown are representative images from 3 independent experiments. **(E)** Quantification of capsule size from India ink staining. Images taken were analyzed with imageJ/Fiji. A minimum of 170 cells of each strain were analyzed. Data are presented as box and whiskers diagram with indicated median and min and max of capsule sized measured for the indicated strain. A mixed effect analysis was performed using the log transformed data. **(F)** Scanning electron microscopy (SEM) analysis of WT and *cbi1*Δ cells in absence and presence of 250 µM BCS.

Compared to many fungal pathogens, *C. neoformans* is relatively tolerant to caspofungin, an antifungal drug that inhibits β-glucan synthesis. However, *C. neoformans* strains with defects in chitosan production are more susceptible to this drug [33]. This observation is consistent with a conserved compensatory increase in cell wall chitin and chitosan among diverse fungi upon treatment with caspofungin [34, 35]. Given the reduced cell wall chito-oligomer content in the copper-starved *cbi1Δ* mutant, we hypothesized that this strain would be more susceptible to caspofungin. Consistent with this hypothesis, we observed a greater than five-fold decrease in the minimal inhibitory concentration (MIC) of caspofungin for the *cbi1*Δ mutant strain compared to the WT and reconstituted strains when incubated in copper-deficient conditions (MIC_50_ *cbi1*Δ 3.1 μM) (Fig 5B, Supp Fig 2G). Together these results suggest that the fungal cell wall mediates durable and adaptive cellular responses to copper availability that affect host cell interactions and antifungal drug activity.

### Depletion of Cbi1 affects several cell wall-associated virulence factors

The *Cryptococcus* cell wall is not only important for stress resistance and modulating immune response but also for several well-established virulence factors such as melaninization and capsule formation. The cryptococcal capsule is composed of highly branching polysaccharides that are covalently attached to components of the outer surface of the cell wall, especially α-1,3 glucans [36]. The thickness of the capsule is induced during incubation conditions that mimic the host environment, including slightly alkaline pH and micronutrient limitation [11]. Strains with mutations in several chitin deacetylase genes, in particular with a *cda2* deletion, typically display enlarged capsules [21]. Similarly, the *cbi1*Δ mutant, in which *CDA2* levels are reduced, produced more surface capsule, especially in copper limiting conditions (Fig 5D-E).

We performed scanning electron microscopic (SEM) analysis of WT and *cbi1*Δ cells after 3d of capsule induction in the absence and presence of 250 μM BCS (Fig 5F) to further analyze the role of Cbi1 in capsule architecture with a higher topographical resolution. Similar to capsule assessment by India ink counterstaining, we observed the most notable alterations in the capsular structure of the *cbi1*Δ strain during copper limitation. Under normal capsule-inducing conditions both strains showed typical capsule architecture with a more dense inner zone and a less dense outer layer with extended capsule fibers [36]. However, when copper deficiency was induced by BCS supplementation, the *cbi1*Δ capsule displayed denser and more interconnected polysaccharide fibers compared to WT, with fewer freely extending individual capsule fibers.

Both copper limitation and cell wall alterations are important for the regulation of melanin, a cell wall-associated antioxidant pigment [23, 29]. Melanin production itself is tightly linked to cellular copper levels since the rate-limiting phenoloxidase enzyme involved in melanin synthesis, laccase-1 (*Cn* Lac1), is functionally dependent on copper [37]. The *cbi1*Δ strain displayed defective melanin production on L-DOPA containing media (Fig 5C). The restoration of normal melanin production by the addition of 5 μM CuSO_4_ is consistent with the known defects in copper acquisition in this strain [5]. We did not observe altered melanin production for the *ctr1*Δ or *ctr4*Δ strains, suggesting that the degree of intracellular copper limitation is greater in the absence of of the Cbi1 protein than in either of the single copper transporter mutants. Also, chitosan is required for the attachment of melanin to the cell surface and strains completely lacking chitosan display a “leaky melanin phenotype” in which melanin diffuses from the cell into the growth medium [21]. Even though depletion of Cbi1 affects cell wall chitosan levels during copper deficiency, we did not observe a leaky melanin phenotype. This is consistent with a reduction but not the complete absence of chitosan in the *cbi1*Δ strain. Together these results suggest that Cbi1-mediated cell wall remodeling and copper homeostasis affect the establishment of several cell wall-associated virulence factors.

## Discussion

In these experiments we demonstrated that copper deficiency, and either deletion of the Cuf1 transcription factor or the copper-binding Cbi1 cell surface protein, renders *C. neoformans* more susceptible to cell wall stress. This central result suggests that copper homeostasis mechanisms affect the integrity of the fungal cell wall. In turn, we demonstrated that genes involved in cell wall chitosan biosynthesis modulate resistance to copper stress. Furthermore, we characterized the biochemical and physiological changes that occur in the cryptococcal cell wall in response to low copper availability. These changes include a reduction in the amount of chitin and chitosan, two structurally related carbohydrates that are typically deeply embedded in the cell wall. Other cell wall-associated processes that were also altered in the *cbi1* mutant include an increased accumulation of surface capsule and a reduction in melanin, two mediators of virulence in this human fungal pathogen. Given the established role of the *Cn* cell surface to mediate immune avoidance, we documented altered interaction of the *cbi1*Δ mutant with macrophages, indicating that these cell wall changes influence the ways in which *Cn* interacts with host innate immune cells.

Copper regulation by the infected host is an important mechanism of immune response to invading pathogens. In contrast to the well-described “nutritional immunity” that primarily involves the sequestering of other essential micronutrients such as iron and zinc away from infecting microorganisms, the host regulation of copper involves a complex coordination of copper sequestion in certain tissues and the induction of very high copper levels in other sites [3, 22, 38, 39]. In this way, microbial pathogens must be able to acquire sufficient copper for cell metabolism and energy production while simultanesously preventing the harmful effects of host-induced copper toxicity. In *C. neoformans*, the Cuf1 transcription factor controls this cellular response to both low and high copper concentrations [2, 4]. In contrast, many other fungal species, both pathogens and nonpathogens, utilize distinct transcriptional regulators for each of these copper states. A genome-wide analysis identified many new and novel copper- and Cuf1-regulated genes, likely reflecting the wide range of copper concentrations encountered by this fungal pathogen in its varied environmental niches [4].

In response to high copper stress, microorganisms have developed multiple means of copper detoxification, including copper efflux systems as well as the induction of the copper-binding metallothienein proteins [2, 39]. In *C. neoformans*, expression of the *CMT1* and *CMT2* metallothionein genes is controlled by Cuf1 in response to elevated copper levels, and these genes are required for fungal survival at sites of high copper exposure such as the host lung tissue [38]. Another reported defense mechanism against the effects caused by high copper stress is the up-regulation of the *ATM1* gene, required for transport of a Fe-S precursor into the cytosol to protect cytosolic Fe-S cluster proteins from mis-metalation caused by unbound free Cu^+^ ions [40]. Recent reports have also demonstrated *Cn* proteomic responses to copper toxicity, including inhibition of protein translation and the induction of ubiquitin-mediated protein degradation [41]. Additionally, metabolic profiling of *Cn* during high copper stress demonstrated large changes in carbohydrate and amino acid metabolites [42]. In this study, we observed that the *CBI1* and *CDA2* genes, which are both regulated by Cuf1, are required for adaptive cell wall remodelling in response to copper stress, specifically by altering the cell wall chitin/chitosan layer.

Despite its toxicity, copper is essential for critical cellular processes, serving as a catalytic cofactor that drives iron uptake and distribution, mitochondrial cytochrome oxidase activity, and reactive oxygen (ROS) detoxification through the Cu/Zn superoxide dismutase (Sod1) [2]. Therefore, in response to low copper stress, microbes have evolved strategies to increase copper uptake efficiency and to direct copper to sites where it is most needed [2, 43, 44]. In *C. neoformans*, increased expression of the copper importer genes *CTR1* and *CTR4* and the *CBI1* gene represents an early response to copper limitation, and is important for fungal survival at sites of copper limitation [4, 5, 22, 30]. A recent study demonstrated another copper sparing mechanism used by *C. neoformans* to adapt to copper limitation in which Cuf1 directs the transcriptional down-regulation of the copper-dependent superoxide dismutase *SOD1* gene and the simultaneous re-localization of the copper-independent Sod2 protein to maintain cellular antioxidant defense levels during copper limitation [45].

Based on our findings we suggest a model in which the fungal cell wall serves two functions relevant to copper homeostasis. First, under conditions of copper excess, cell wall carbohydrates bind copper ions to prevent cytotoxic copper stress. Second, under conditions of copper deficiency, the cell wall releases bound copper to metal transporters for maintenance of cell homeostasis. Our data also suggest that the Cbi1 protein is one component of this copper acquisition process between the Ctr1 copper transporter and the fungal cell wall. *CBI1* and *CTR1* are among the *Cn* genes with the highest degree of regulation by the copper-sensing Cuf1 transcription factor in conditions of Cu depletion [4, 5]. The Cbi1 and Ctr1 proteins interact both physically and genetically [5]. Accordingly, the *cbi1*Δ and *ctr1*Δ mutants display some similar phenotypes: poor growth in the presence of copper deprivation and cell wall stress.

However, the Cbi1 protein appears to have functions that are independent of simply participating in Ctr1-mediated copper transport into the cell. The *cbi1*Δ strain has a more severe, but copper-remediable, melanin production defect than the *ctr1*Δ strain, suggesting a greater degree of intracellular copper limitation or altered copper acquisition. Moreover, only the *cbi1*Δ mutant, and not the *ctr1*Δ or *ctr4*Δ copper transporter mutants, displayed enhanced resistance to toxic copper levels. These results begin to elucidate a new role for cell wall polysaccharides in adaptation to environmental changes. Since cell wall polysaccharides such as chitin and chitosan are known to avidly bind divalent metal ions such as Cu^2+^, it is likely that their presence in the fungal cell wall serves to similarly bind copper encountered in the environment. At lower levels of copper exposure, cell wall-associated copper might provide a readily accessible storage site for the copper transporter complexes. At higher levels of copper, polymers such as chitosan might bind copper, protecting the cell body and membranes against the toxic effects of free Cu^2+^ ions.

The Cbi1 protein displays some degree of sequence similarity to proteins such as LPMOs that bind complex carbohydrate surfaces and promote further structural alterations by specific hydrolases [46]. LPMOs have been best explored in the context of degradation of crystalline cellulose and chitin [47]. We have previously demonstrated that Cbi1 binds copper, but it lacks the redox activity that defines the LPMO class of enzymes [5, 6]. Its copper-binding activity, and its physical association with the Ctr1 protein, suggest that Cbi1 might act as an intermediary to shuttle copper from the cell wall storage sites to the copper importer system. Our TEM data here support this model. In these images the electron density of the chitin/chitosan cell wall layer remains unchanged in the *cbi1*Δ mutant during extracellular copper starvation. In contrast, the electron density of this layer in WT cells decreases during copper starvation, consistent with enhanced cellular import of this metal ion mediated in part by Cbi1.

Here we have demonstrated that the absence of Cbi1 is also associated with transcriptional changes in cell wall genes, and that these transcriptional changes result in functional consequences for the *Cn* cell wall. These changes include reduced levels of chitin and chitosan as well as altered cell wall architecture, especially in the presence of copper limitation. As a result, cell wall-associated virulence factors are altered in function, resulting in changes in the interaction with host immune cells. The specific activity of the Cbi1 protein has yet to be determined. Also, it is not yet clear whether the cell wall changes in the *cbi1*Δ strain are directly related to Cbi1 function or whether they represent compensatory cellular changes in response to stress. This type of cell wall adaptation to cell stress is commonly observed in other conditions [48]. For example, cell wall chitin levels are increased in the human fungal pathogens *Aspergillus fumigatus* and *C. neoformans* during treatment with the beta-glucan synthase inhibitor capofungin [33]. Blunting this adaptive cell wall chitin response renders these cell more susceptible to the activity of this antifungal agent. Similarly, we demonstrated here that the *cbi1*Δ mutation, and its downstream defective chitin response, results in a similar degree of enhanced caspofungin susceptibility as mutation in chitin synthesis genes themselves.

Our observations of significant reductions in cell wall chitosan in the copper-starved *cbi1*Δ strain also suggest that this cell wall polymer may contribute to copper homeostasis. In contrast to many ascomycete fungal pathogens, *C. neoformans* contains much more chitosan in the inner chito-oligomer layers of the cell wall [8]. Chitosan is a relatively de-acetylated form of chitin. Both chitin and chitosan exist in varying sizes, the polymer length determined by both endogenous and exogenous synthases and chitinases. In fact, the size of the chitin and chitosan molecules, as well as their relative degrees of acetylation/deacetylation, determine their immunogenicity. Therefore, both host and microbe possess intricate means to regulate chito-oligomer molecular size and acetylation status [8, 13, 49].

In *C. neoformans*, four putative chitin deacetylases contribute to the conversion of chitin to chitosan: Cda1, Cda2, Cda3, and Fpd1. Prior mutational analysis revealed that a *Cn cda1,2,3* triple mutant is devoid of most measurable cell wall chitosan during vegetative growth [21]. Although the Fpd1 enzyme is not required for chitosan conversion from chitin under normal growth conditions, it may contribute to the further deacetylation of pre-formed chitosan [50]. Consistent with the decrease of cell wall chitosan in the *cbi1*Δ mutant, we identified two of the four chitin deacetylase genes, *cda2* and *fpd1*, to be down-regulated in copper-deficient *cbi1Δ* cells. Hence, Cda2 and Fpd1 may be involved in regulating cell wall chitosan levels in response to cellular copper levels. Notably, *cda2* was previously identified as a Cuf1-regulated gene [4], which further strengthens the connection between cell wall chitosan and copper homeostasis. Therefore, Fpd1 together with Cda2 could potentially be involved in modulating the deacetylation ratio of chitosan molecules, and by doing so modulating the copper binding capacity of the aggregate cell wall sugar polymers in response to changing copper levels.

Cell wall chitin, among other cell wall sugars, is recognized by host pattern recognition receptors triggering immune activation and defense mechanism. Therefore, masking chitin is one important tool to evade immune defense. One strategy for dampening the immune recognition of chitin is its deacetylation to form chitosan [49]. Our cell wall analysis revealed a complex set of changes in response to copper homeostasis and the *cbi1*Δ mutation. Although the total levels of chitin and chitosan were decreased in the *cbi1*Δ strain, the degree of exposure of these cell wall sugars was greater in this strain compared to WT. Similar alterations in chitin/chitosan exposure have been shown to fundamentally alter the degree of immunological masking of fungal cells from recognition by host immune cells. In fact, there was a striking correlation between the degree of exposed chitin/chitosan exposure (as measured by WGA Alexa488 staining) and the activation of a macrophage TNF-α response in fungal co-culture [13, 32]. Also, lectins and monocolonal antibodies that block chitin recognition have been proposed as adjunctive therapeutic strategies for cryptococcosis and other fungal infections [51].

In summary, we have defined cell wall changes that mediate adaptation of a fungal pathogen during conditions associated with human infection, including both high and low copper levels. In this way, the cell wall serves as both storage site for copper during low copper levels, as well as a copper-binding organelle to prevent excessive intracellular accumulation during copper toxicity. We have further defined cellular roles for a unique copper-binding protein that serves to mediate copper transfer between the fungal cell wall and copper import proteins.

## Material and Methods

### Strains, media and growth conditions

*Cryptococcus neoformans* strains used in this study are shown in **Supp Table 1.** All strains were generated in the *C. neoformans var. grubii H99* background. For strain creation, DNA was introduced into *C. neoformans* by biolistic transformation [52]. Yeast extract (2%)-peptone (1%)-dextrose (2%) (YPD) medium supplemented with 2% agar and 100 μg ml-1 of nourseothricin (NAT), 200 μg ml-1 of neomycin (G418) or 200 μg ml-1 of hygromycin B (HYG) was used for colony selection after biolistic transformation. Cloning strategies as well as plasmids and oligos used for creation of *Cryptococcus* transformation contructs are described in **Supp Table 2-3.** Transformants were screened by PCR and Southern blot for intended mutations. Cbi1-HA expression among relevant transformants was confirmed by western blot.

Strains were cultivated in either synthetic complete (SC) medium (MP Biomedicals) or YPD at 30°C. To induce Cu sufficiency or deficiency, media was supplemented with indicated concentrations of CuSO_4_ or the Cu^+^ chelator bathocuproine disulfonate (BCS), respectively. Alternatively, to BCS supplementation, strains were cultivated in Yeast extract-peptone medium supplemented with 3% Glycerol and 2% Ethanol (YPEG). For galactose-regulated expression induction, SC+2% Galactose (SC-Gal) or SC+2% Glucose (SC-Glu) was used and supplemented as indicated. To analyze cell wall associated phenotypes, caffeine (0.5 mg/ mL), NaCl (1.5 M), SDS (0.01%), Congo red (0.5%) and Calcofluor White (1.5 mg/mL) were added to SC medium supplemented with CuSO_4_ or BCS as indicated. For growth phenotype analysis on solid medium plates, a 6-fold serial dilution, starting at OD_600_ 0.25, of strains was spotted and incubated for indicated time and temperature. For assessment of melanization, overnight cultures in YPD were washed once in PBS and resuspended in PBS to OD_600_ 2.5. Next, 5 to 10 μL of the resuspended culture were spotted onto L-3,4-dihydroxyphenylalanine (L-DOPA) media (7.6 mM L-asparagine monohydrate, 5.6 mM glucose, 22 mM KH2PO4, 1 mM MgSO4.7H2O, 0.5 mM L-DOPA, 0.3 μM thiamine-HCl, 20 nM biotin, pH 5.6). L-DOPA plates were incubated at 30°C for 2 days. To induce capsule, strains were incubated in CO2-independent tissue culture medium supplemented as indicated (TC, Gibco) for 72 hours with shaking at 37°C, followed by staining with India Ink or fixation for scanning electron microscopy (SEM).

### RNA isolation and qRT-PCR

For *ROM2* transcript analysis *C. neoformans* overnight cultures grown in synthetic complete (SC) medium (MP Biomedicals) were diluted to OD_600_ 0.3, and cultures were supplemented and cultivated as indicated. For cell wall synthesis genes transcript analysis and copper status analysis, *C. neoformans* overnight cultures grown in YPD medium were diluted to OD_600_ 0.05, supplemented as indicated and cultivated for 24h at 30°C. For RNA extraction, cells were cultivated as indicated. Cultures were harvested, washed 1x with PBS and flash frozen on dry ice, followed by lyophilization. RNA was extracted using the RNeasy Plant Mini Kit (Qiagen) with optional on-column DNAse digestion. cDNA for real time-PCR (RT-PCR) was prepared using the Iscript cDNA synthesis kit (Biorad). For RT-PCR, cDNA was diluted 1:5 in RNase-free water, added to ITAQ Universal SYBR Green Supermix (Bio-Rad) per protocol instructions and analyzed on a CFX384 C1000 ThermoCycler (BioRad, ROM2 analysis and Cu status analysis). For analysis of cell wall transcripts, the diluted RNA was mixed with the PowerUP SYBR Green Master mix (applied biosystems) per protocol instruction and analyzed on a QuantStudio 6 Flex (applied biosystems). Oligos used for qRT-PCR analysis are shown in **Supp Table 3.** C_T_ values were determined using the included CFX Maestro software (BioRad) or the QuantStudio 6 Flex, respectively. Gene expression values were normalized to the housekeeping gene *GAPDH* and expression fold changes determined by the ΔΔC_T_ method. For all qRT-PCR studies, a minimum of 3 independent biological replicates were used for the analysis of mRNA expression changes.

### Inductively coupled plasma mass spectrometry (ICP-MS)

Cell associated metals (Fe and Cu) were quantified from lyophilized yeast. In short, yeast cells were treated as indicated, spun down and washed 2x with ICP-MS grade water. In the last wash step, cell were counted, spun down and lyophilized. Samples were digested in 300 uL 50% ICP-MS grade Nitric Acid for 1h, at 90°C and cooled down overnight at RT. Metal content was analyzed by ICP-MS at the Oregon Health Sciences University elemental analysis facility on an Agilent 7700X ICP-MS.

### Liquid growth assays

All liquid growth analysis were performed in 96 well plates. For the growth analysis in YPEG in the presence of the surface stressor SDS, overnight cultures (YPD, 30°C), were harvested and washed 1x with YPEG and then normalized to OD_600_ 2.0 (in YPEG). Growth media was supplemented as indicated and filled into 96 well plate (195 µL each well). Wells were inoculated with 5 μL of indicated strain (final OD_600_ =0.05). Plates were covered with a semipermeable membrane (Breathe-Easy, Diversified Biotech) and incubated at 30°C with shaking at 1150 rpm in a Finstruments shaker instrument. Growth graphs of the indicated strains at the conditions analyzed were generated by plotting the OD_600_ readings normalized to WT-YPEG growth at the 24 h time point. Three biological replicates were performed.

For minimal inhibitory concentration analysis (MIC), overnight cultures (SC, 30°C), were harvested and washed 1x with PBS and then set to OD_600_ 0.25 (in PBS) and stored on ice until further usage. 2x concentrated working stocks of caspofungin were prepared in SC medium (final concentration in assay ranged 100 to 0.78 μg/mL). Cells were diluted 1:100 in either SC or SC+200 μM BCS (final concentration in assay 100 μM BCS). In 96 well plate, 100 μL of diluted cells were mixed with 100 μL of 2x concentrated caspofungin stocks. The plate was covered with a semipermeable membrane (Breathe-Easy, Diversified Biotech) and incubated at 30°C with shaking at 1150 rpm in a Finstruments shaker instrument for 24h. Growth graphs of the indicated strains at the conditions analyzed were generated by calculating the relative growth of the drug-treated condition in relation to the untreated condition (drug-treated OD600/untreated OD600). Four biological replicates were performed.

### Transmission electron microscopy (TEM)

Overnight cultures (YPD, 30°C) were harvested and washed 1x with YPD. Indicated strains were inoculated to an OD_600_ of 0.05 in 50 mL YPD + 10 μM CuSO_4_ (=Cu sufficiency) or 250 μM BCS (=Cu deficiency) and cultivated for 24h at 30°C. In the following day, 50 to 100 μL of the culture were harvested and washed 1x with PBS. Next, cells were pelleted and overlayed with fixative (4% formaldehyde, 2% Glutaraldehyde in PBS) and incubated for 4h at RT. Then, fixative was removed, and the sample was washed twice with 1x PBS, with a 10 minute incubation time at RT in between wash steps. After last washing step, the PBS was removed and 1% OsO_4_ was added to the sample to complete cover. The tube is sealed and incubated for 1h at RT in the dark. Then the OsO_4_ was removed, and the sample rinsed with 1x PBS, 2 times for at least 10 minutes each time (RT). After last PBS rinse, residual PBS was removed and sample was rinse with 0.1N acetate buffer, 1 time at least 10 minutes each time (RT). The acetate buffer was removed, and the sample was stained with 0.5% uranyl acetate (UA) for one hour, RT. Once staining is complete, the uranyl acetate was removed and the sample rinsed with 0.1N acetate buffer, 2 times at least 10 minutes each time. In the next steps the samples were dehydrated in several ethanol incubation steps by rinsing twice, at least 10 minutes each time, with 30% Ethanol, 50% Ethanol, 70% Ethanol and 90% Ethanol. Finally, the sample were rinsed 3 times, at least 10 minutes each time, with 100% Ethanol.

Once the dehydration was complete, ethanol was removed and the dehydrated sample was embedded into resin (53,5% (w/v) resin, 20.5% (w/v) DDSA, 26% (w/v) NMA, 1.4% (v/v) DMP-30). The sample were incubated in resin mix at RT overnight. The following day, samples were incubated at 50-60°C for 10 minutes, the old resin mix was replaced by freshly made resin mix and incubated for 10 mins at RT, followed by 10 minutes at at 50-60°C. This resin wash step was repeated one more time, followed by a 48h incubation at 50-60°C. The embedded samples were cut into 70nm thick sections on an Ultracut microtome and placed on TEM grids. The sections were counterstained with uranyl acetate and lead citrate and then imaged on an FEI Technai G2 Twin transmission electron microscope. Cell wall thickness was measured using ImageJ (Cu sufficient WT: 8 cells, Cu deficient WT: 7 cells, Cu sufficient *cbi1Δ:* 14 cells and Cu deficient *cbi1Δ:* 10 cells). The staining contrast in the cell wall was measured using Image J gray scale measurement tool.

### Scanning electron microscopy (SEM)

Overnight cultures (YPD, 30°C) were harvested and washed 1x with PBS. Indicated strains were inoculated to an OD_600_ of 0.1 in 25 mL CO_2_-independent medium (Gibco) or 25 mL CO_2_-independent medium supplemented with 250 μM BCS and cultivated for 3d at 37°C. 5 mL of each culture were harvested, checked for capsule formation by India ink stain and washed 3x with PBS (without calcium and magnesium). Cells were fixed for 1h at RT, using 2.5% glutaraldehyde in PBS and washed 3 times with PBS and checked for intact capsule by india ink staining. Then, cells were mounted onto poly-L-lysine-coated coverslips (Neuvitro, 12mm, #1 thickness coverlsips) and incubated for 20 min at RT. After mounting, cells were sequentially dehydrated in several ethanol washes (1x 30%, 1X 50%, 1X 70%, each 5 min RT, followed by 1x 95% and 2x 100%, 10 min RT). After dehydration mounted cells were stored in 100% ethanol until the critical point drying. Cell samples were critical point dried with a Tousimis 931 critical point dryer (Rockville, Maryland) and coated with gold-palladium using a Cressington 108 sputter-coater (Watford, United Kingdom). Samples were mounted and imaged on a Hitachi S-4700 scanning electron microscope (Tokyo, Japan).

### Cell wall isolation and analysis

Overnight cultures (YPD, 30°C) were harvested and washed 1x with YPD. Indicated strains were inoculated to an OD_600_ of 0.05 in 50 mL YPD + 10 μM CuSO_4_ (=Cu sufficiency) or 250 μM BCS (=Cu deficiency) and cultivated for 24h at 30°C. The following day, 10 to 25 mL of the cells were harvested and washed twice with dH_2_O. In the last wash step, cells were counted, spun down and lyophilized. Chitin and chitosan levels were quantified from lyophilized yeast using a modified MBTH (3-methyl-benzothiazolinone hydrazine hydrochloride) method as previously described [13]. β-glucan was quantified using the megazyme yeast β-glucan kit. In short lyophilized yeast were milled using glass beads, resuspended in 800 µL 2 M KOH and transferred into a new 12 ml reaction tube and stirred for 30 mins in an ice water bath. Then, 3.2 mL of 1.2 M sodium acetate pH 8.3 and 40 µL glucazyme was added and the sample stirred for 2 min. The sample was transferred to a 15 mL screw cap tube and incubated ON at 40°C in a water bath. The next day, 10 mL dH_2_O was added to the samples, mixed thoroughly and centrifuged for 10 mins at 3000 rpm. Then, 100 µL of the supernatant was mixed with GOPOD reagent and incubated for 20 mins at 40°C in a water bath. 2 x200 µL of each sample were transferred into a 96 well plate and read (against reagent blank) at 510 nm. A standard curve was prepared using the manufacturer’s supplied D-glucose standard solution and mg glucose was calculated using equation provided by manufacturer. Measured values were normalized by cell count.

### Cell wall staining and flow cytometry

Prior to analysis cells were treated as indicated. To visualize chitin, cells were harvested and stained with 100 μg/ml Alexa488-conjugated wheat germ agglutinin (WGA, Molecular Probes) for 35 minutes in the dark, RT, followed by 25 μg/ml calcofluor white (CFW, Fluka Analytical) for 10 minutes, RT. After staining, cells were washed 2x with PBS and were resuspended in 20-50 µL PBS for microscopic analysis. Alexa488-WGA was imaged using a GFP filter and CFW was imaged using a DAPI filter. For flow cytometry, cells were Alexa488-WGA and CFW stained as previously described, washed 2x with PBS and set to 10^6^ cells (in 1 mL PBS). Alexa488-WGA stained cells were analyzed using a 488 nm laser and CFW cells were analyzed using a 405 nm laser. The FACS analysis was performed at the Duke Cancer Institute Flow Cytometry Shared Resource using a BD FACSCanto II flow cytometer. Data was analyzed using FlowJo v10.1 software (FlowJo, LLC). For analysis only single cells were used (gated using the FSC/SSC plot). For chitin exposure analysis cells were gated in the CFW intensity/ Alexa488-WGA intensity scatter plot. Additional histograms with mean fluorescence intensity (MFI) on the x-axis and cell counts on the y-axis were created. Unstained cells were used as negative controls.

To visualize chitosan, cells were treated as indicated, harvested and washed 2x with McIlvaine’s buffer (0.2 M Na2HPO4, 0.1 M citric acid, pH 6.0). Then, cells were stained using 500 µL of 300 μg/ml Eosin Y in McIlvaine’s buffer for 10 minutes at room temperature in the dark. Cells were then washed 2x with McIlvaine’s buffer and resuspended in 20-50 µL McIlvaine’s buffer. Cells were visualized using a GFP filter.

### Microscopic quantification

Differential interference microscopy (DIC) and fluorescent images were visualized with a Zeiss Axio Imager fluorescence microscope (64X objectives). Images were taken with an AxioCam MRm digital camera with ZEN Pro software (Zeiss). The same exposure time was used to image all strains analyzed. Images were analyzed using ImageJ/Fiji software. Gray scale values were measured and normalized towards cell count. The intensity of the control strain (=Cu sufficient WT) was set to 1. Results are reported as relative fluorescence intensity +/-standard error of the means.

Cells sizes were measured using the ImageJ measurement tool. Capsule thickness was calculated using the equation:

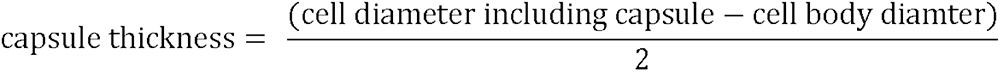

### Generation of bone marrow derived macrophages

Murine bone marrow cells were isolated from A/J mice and prepared as previously described [31]. Briefly, femurs and tibias were isolated from mice. Each bone was flushed with 5 to 10 ml cold PBS using a 27½ gauge needle. Red blood cells were lysed in 1x RBC lysis buffer (0.15 M NH_4_Cl, 1 mM NaHCO_3_, pH 7.4) and cells were resuspended in 1x Dulbecco’s modified Eagle’s medium (DMEM; + 4.5 g/L D-Glucose, + L-Glutamine, +110 mg/L sodium pyruvate) with 1 U/ml pencillin/streptomycin. Bone marrow cells were cryopreserved in 90% FBS/10% endotoxin-free DMSO at a concentration of 1 x 10^7^ cells/ml.

BMMs were differentiated in BMM medium (1x Dulbecco’s modified Eagle’s medium [DMEM; + 4.5 g/L D-Glucose, + L-Glutamine, +110 mg/L sodium pyruvate], 10% fetal bovine serum [FBS; non-heat inactivated], 1 U/ml penicillin/streptomycin) with 3 ng/ml recombinant mouse GM-CSF (rGM-CSF; R&D Systems or BioLegend)) at a concentration of 2.5 x 10^5^ cells/ml in 150 x 15 mm petri plates at 37°C with 5% CO_2_. The media was refreshed every 3–4 days and the cells were harvested after ∼7d or when confluency was achieved. The Duke University Institutional Animal Care and Use Committee reviewed and approved the protocol for the macrophage harvesting. Protocol registry number A102-20-05.

### Macrophage co-incubation and TNF-α quantification

Prior to co-incubation, overnight cultures of *C. neoformans* strains (YPD, 30°C) were harvested and washed 1x with YPD. Indicated strains were inoculated to an OD_600_ of 0.05 in 50 mL YPD + 10 μM CuSO_4_ (=Cu sufficiency) or 250 μM BCS (=Cu deficiency) and cultivated for 24h at 30°C. To prepare BMMs for the co-incubation assay, BMMs were counted (by hemocytometer, with Trypan blue to discount dead cells), plated in BMM medium in 96-well plates at a concentration of 5 x 10^4^ cells/well and incubated at 37°C with 5% CO_2_ overnight. The next day C. *neoformans* cells were washed 2x with PBS, counted, and added to BMMs containing 96-well plates at a concentration of 5 x 10^5^ fungal cells per well (10:1 *C*. *neoformans* cells:BMMs). Co-cultures were incubated for 6h at 37°C with 5% CO_2_. Supernatants were collected and stored at -80°C until analysis. Secreted TNF-α was quantified in supernatants by enzyme-linked immunosorbent assay (ELISA; BioLegend).

### Statistical analysis

For all data error bars represent statistical errors of the means (SEM) of results from a number of biological replicates (N), as indicated in figure legends. Before statistical analysis was conducted, data from all experiments was log transformed for comparison of proportions. Statistical analysis was performed with GraphPad Prism software v9. The statistical tests chosen for each experiment and their results (i.e., p values) are indicated in figure legends. Asterisks in figures correspond to statistical significance as follows: ****, P < 0.0001; ***, P = 0.0001 to P < 0.001; **, P = 0.001 to P < 0.01; *, P = 0.01 to P < 0.05; ns (not significant), P > 0.05.

## Acknowledgements

We thank the Duke Cancer Institute for the use of the Flow Cytometry Shared Resource. The Transmission electron microscopy was performed in part at the Duke University Shared Materials Instrumentation Facility (SMIF), a member of the North Carolina Research Triangle Nanotechnology Network (RTNN), which is supported by the National Science Foundation (award number ECCS-2025064) as part of the National Nanotechnology Coordinated Infrastructure (NNCI). Scanning electron microscopy was performed at the Chapel Hill Analytical and Nanofabrication Laboratory, CHANL, a member of the North Carolina Research Triangle Nanotechnology Network, RTNN, which is supported by the National Science Foundation, Grant ECCS-1542015, as part of the National Nanotechnology Coordinated Infrastructure, NNCI. ICP-MS measurements were performed in the OHSU Elemental Analysis Core. We thank Dr. Martina Ralle for her help and insight in sample preparation and ICP-MS data acquisition and Dr. Aaron D. Smith for insightful discussions of the data and proposed model. This work was also supported by funding from the National Institute Health: NIAID R01 AI074677 (JAA) and NIGMS R01GM041840 (DJT and JAA) and the German Research Foundation grant PR 1727/1-1 (given to CP).

## Figure Legends

**Supp Fig. 1**: **(A)** Growth analysis in presence of cell wall/ surface stressors. The spotting assay was performed on SC supplemented with indicated amounts of cell wall and cell surface stressors. Indicated strains were grown overnight in SC at 30°C. Cells were diluted to OD_600_ of 0.25 and a serial 1:10 dilution was spotted on to media plates. Plates were incubated at 30°C for 2-4d. This figure shows a representative image from 3 independent spotting experiments. **(B)** Growth rate of copper sufficient or deficient WT and *cbi1Δ* cells. Cells were incubated in YPD supplemented with 10 µM CuSO_4_ (Cu sufficiency) or with 250 µM BCS (Cu deficiency) for 24h at 30°C. Growth was measured through 0D_600_. Presented is the average +/-SEM of 5 biological replicates. **(C)** Colony forming unit (CFU) analysis of copper sufficient or deficient WT and *cbi1Δ* cells. Cells were treated as described in (B). After 24h of growth, cells were diluted to OD_600_ 1. 200 μL of a serial 1:1000 dilution were plated onto YPD plates and colonies were counted after 3d of incubation at 30°C. The CFU of copper sufficient WT was set to 100%. Presented is the average +/-SEM of the relative CFU (as compared to copper sufficient WT) from 4 biological replicates. **(D)** qRT-PCR analysis using *CMT1* and *CTR4* as indicator for Cu toxicity or deficiency. Indicated cells were cultivated as described in (B) and used for RNA extraction, followed by cDNA synthesis. Presented is ΔΔC_T_ of copper deficiency: copper sufficiency. The average +/-SEM from 3 biological replicates is shown. **(E-F)** ICP-MS based metal quantification of cell associated copper (E) and Iron (F) in pg metal per 10^6^ cells. Indicated strains were grown as described in (B). Presented is the average +/-SEM from 3 biological replicates.

**Supp Fig. 2**: **(A)** β-glucan quantification of copper sufficient and copper deficient wt, *cbi1*Δ and *cbi1*Δ*^C-WT^* complemented cells. Strains were incubated for 24h in YPD+ 10 uM CuSO4 (Cu sufficiency) or YPD +250 uM BCS (Cu deficiency) and then harvested, cell counted and lyophilized. The Megazyme yeast b-glucan kit was used for quantification of b-glucan from lyophilized cells. Values are shown in ug Glucose / 10^7^ cells. Presented is the average +/-SEM of 3 biological replicates. **(B)** Calcofluor white (CFW) and wheat germ agglutin (WGA)-Alexa 488 staining for chitin of copper sufficient or deficient WT, *cbi1*Δ and and Cbi1 WT complemented *cbi1*Δ (*cbi1*Δ*^C-^*^WT^) cells. Strains were cultivated as described in (A). Shown is the mean +/-SEM of the relative CFW intensity from 3 independent experiments. The CFW intensities were measured with ImageJ/Fiji and normalized to cell count. Shown is the relative CFW intensity (copper sufficient WT set to 1). A 1-way ANOVA was performed using GraphPad Prism from log transformed data. **(C)** EosinY staining for chitosan of copper sufficient and deficient WT, *cbi1*Δ and *cbi1*Δ*^C-^*^WT^ complemented cells. Strains were cultivated as described in (A), followed by EosinY staining. Shown are representative images for 2 two independent experiments. Five independent treatments and stainings were performed. **(D)** Relative EosinY intensity from 5 independent experiments. The EosinY intensities were measure with ImageJ/Fiji and normalized to cell count. Shown is mean +/-SEM of the relative EosinY intensity (copper sufficient WT set to 1). A 1-way ANOVA was performed using GraphPad Prism from log transformed data. **(E-F)** FACS analysis of CFW and WGA-Alexa 488 stained cells. WT, *cbi1*Δ and *cbi1*Δ*^C-WT^* complemented cells were cultivated as described in (A) **(E)** WGA-Alexa 488 staining histogram representation of the FACS analysis depicted in Fig 3. **(F)** CFW-staining histogram representation of FACS analysis depicted in Fig3. **(G)** Minimal inhibitory concentration (MIC) analysis of Caspofungin during copper sufficiency. WT, *cbi1*Δ and *cbi1*Δ*^C-WT^* complemented cells were grown in 96-well liquid cultures in SC media supplemented with 0 to 100 ug/mL Caspofungin. OD_600_ were measured after 24h growth at 30°C and the OD_600_ of the non-treated condition was set to 1. Presented is the mean +/-SEM of 4 independent experiments.

